# Context-dependent toxicity of human Tau isoforms in a *Drosophila* tauopathy model

**DOI:** 10.64898/2026.03.20.713147

**Authors:** Yelena Ivanova, Miguel Ramirez-Moreno, Jie Liu, Leila Abtahi, Baidong Wu, Amber S. Cooper, Zengbo Wang, Douglas W. Allan, Amritpal Mudher, Aaron A. Comeault, Lovesha Sivanatharajah

## Abstract

Tauopathies are characterised by progressive deterioration of brain regions due to abnormal accumulation of the microtubule-associated protein tau (MAPT). Alternative splicing of MAPT pre-mRNA results in six tau isoforms, which are classified into two groups depending on the number of microtubule-binding domain repeats (3R vs 4R). Although many tauopathies are 3R or 4R-specific, the relative contributions of individual isoforms to neurotoxicity remain incompletely understood. To systematically characterise differences in tau isoform toxicity, we created a novel set of *Drosophila* lines expressing equivalent amounts of the six human tau isoforms (hTau) at levels sufficient to induce visible phenotypes. Using a variety of assays including survival, negative geotaxis and tissue-level or cell-type-specific degeneration, we found that hTau isoform toxicity is not uniform across different biological contexts. Despite generally higher toxicity of 4R isoforms compared to 3R, the effects of individual hTau isoforms varied with the temporal window of expression, tissue type, and neuronal identity. Restricting hTau expression to small homogeneous neuronal populations enabled detailed analysis of isoform-specific degeneration. Neurons previously observed to be vulnerable or resilient to hTau toxicity exhibited differences in the onset and progression of degeneration, suggesting that resilience may be an early and transitory state, with most or all neurons eventually succumbing to tau toxicity over time. Notably, these differences in toxicity were not readily explained by variations in hTau abundance and phosphorylation. Together, our findings demonstrate that tau toxicity is highly context-dependent, clearly isoform-specific, and shaped by interactions between tau and its cellular environment.

## Introduction

The Microtubule-associated Protein Tau (MAPT) is a neuronally expressed protein that plays an important role in maintaining neuronal stability and facilitating axonal transport [1]. Abnormal tau accumulation is characteristic of several neurodegenerative diseases including Alzheimer’s Disease (AD), that are collectively termed tauopathies. In these diseases, tau becomes hyperphosphorylated, which reduces its affinity for microtubules and leads to cytoskeletal destabilisation [2]. Pathological tau forms oligomers and larger aggregates (e.g., neurofibrillary tangles, NFTs), which can disrupt normal cellular transport and synaptic function, ultimately contributing to neuronal degeneration and death [3].

In the adult human brain there are six major tau isoforms, all arising from the alternative splicing of the MAPT pre-mRNA [4]. Exons 2, 3, and 10 are alternatively spliced, with exon 3 never appearing independently of exon 2. This translates to differences in the number of N-terminal motifs (0-2N, exons 2 and 3) and microtubule-binding repeats (3-4R, exon 10). The smallest tau isoform, 0N3R, is 48 kDa, whilst the largest isoform, 2N4R, is 67 kDa. Depending on exon 10 inclusion in the mRNA, the isoforms can be divided into 3R and 4R groups. During development, hTau expression is restricted to 3R isoforms, with 0N3R being the only isoform found in the foetal brain, whereas both 3R and 4R hTau isoforms are expressed in the adult [4], [5].

In healthy adult brains, the levels of 3R and 4R isoforms are approximately equal, and an imbalance in the 3R:4R ratio is associated with disease [6]. In AD, both 3R and 4R isoforms are found within NFTs [4], but many tauopathies are 3R- or 4R-specific, with different types of tau isoforms prevalent in the aggregates. In Pick’s Disease there is a predominance of 3R tau, while in Progressive Supranuclear Palsy (PSP), Corticobasal Degeneration (CBD), Argyrophilic Grain Disease (AGD), and Globular Glial Tauopathy (GGT) there is a predominance of 4R tau [7] [8]. These disorders also differ in the cellular and regional distribution of pathology, as well as in their clinical presentation, supporting the idea that specific neuronal populations are selectively vulnerable to distinct tau isoforms [9]. This suggests that the unique molecular environment of a neuron type may shape its vulnerability to either 3R or 4R species, ultimately contributing to the divergent neuropathological landscapes observed across these diseases.

To date, *Drosophila* disease models have offered a fast and highly translatable experimental platform extensively used to study human tau (hTau) isoforms and their effects on neurodegeneration (reviewed in [10], [11]). Comparisons of the hTau isoforms 0N3R and 0N4R in *Drosophila* have found differences in isoform toxicity [12]. Where 0N3R expression was shown to result in axonal transport, locomotor defects, and shorter lifespan, 0N4R expression led to more severe neurodegeneration and memory impairment. A separate study further demonstrated differences in neurotoxicity between the 3R and 4R isoforms [13]. 0N4R and 2N4R expression induced a rough eye phenotype indicative of neurodegeneration, whereas 0N3R did not. Furthermore, this study suggested that developmental expression of 4R hTau was required for the emergence of adult phenotypes like photoreceptor loss and reduced fly lifespan [13].

Collectively, these studies highlight differences in the neurotoxic effects of 3R and 4R hTau isoforms in *Drosophila*, but the contributions of individual isoforms to tau pathology remain understudied. In particular, 1N3R and 1N4R isoforms are often excluded from the analysis of tau effects on neurodegeneration, limiting our understanding of the full profile of tau toxicity. Furthermore, until recently, many *Drosophila* studies have not systematically accounted for genetic variability like the one arising from the genomic insertion site of transgenic tools (reviewed in [10]). Transgenes are subject to position-effect variegation, leading to potential differences in isoform expression levels that complicate comparisons of isoform-specific toxicity. For studies aimed at understanding isoform specific differences in Tau toxicity, it is therefore imperative to use models with equivalent expression of all isoforms being assessed.

Previously, Fernius *et al.* (2017) created a set of transgenic fly lines where the six hTau isoforms were inserted into the same genomic locus, VK00018, to enable direct comparison of isoform-specific effects [14]. Analyses of fly survival and behaviour found an overall similar isoform toxicity, with 2N3R expression leading to a significantly reduced lifespan compared to 2N4R. Further investigation of these hTau isoforms in another study found similar effects on oxidative stress, locomotor behaviour, and survival [15]. In both studies, the effects of the six hTau isoforms on the phenotypes assayed were very mild. One possible explanation is that the hTau expression levels in these lines did not reach the threshold required to induce distinct degenerative phenotypes.

To account for all these challenges, we generated a new set of *Drosophila* lines showing robust expression of hTau isoforms. The six UAS-hTau constructs were inserted into the same landing site, VK00037, but in contrast to *Fernius et al. (2017)*, 10xUAS constructs were used to greatly boost expression of all the hTau isoforms. Examination of these new lines for classic tau-dependent phenotypes found very clear differences in isoform toxicity. Across all examined tissues, we observed a general trend in which 4R isoforms showed greater toxicity than 3R isoforms. However, the precise hierarchy of isoform-induced toxicity differed between the tissues, indicating that tau-induced degeneration is highly tissue-dependent and shaped by the interactions between specific isoforms and the cell-type in which they are expressed, including its unique local molecular landscape. Examination of neurons previously determined to be vulnerable or resilient to hTau0N3R toxicity [16] revealed differences in the onset and progression of degeneration, with neurons classed as resilient exhibiting degeneration at later stages. This suggests that resilience may be an early and transitory phase, with neurons succumbing with time to the toxicity caused by any tau isoform.

## Results

### New Drosophila transgenic lines express hTau isoforms at similar levels

DNA sequences for each of the six hTau isoforms (Fig. 1A) were placed downstream of ten GAL4 upstream activation sequences (10xUAS). These constructs were flanked by two gypsy insulators and inserted into the same landing site, VK00037, on chromosome 2 (Fig. 1B). To determine whether hTau isoforms were expressed at comparable levels in adult *Drosophila* central nervous system (CNS), we used the TARGET system, combining the pan-neuronal elav-GAL4 driver with the ubiquitously expressed temperature-sensitive Gal80 repressor (Gal80^ts^) to control transgene induction [17].

**Figure 1.**
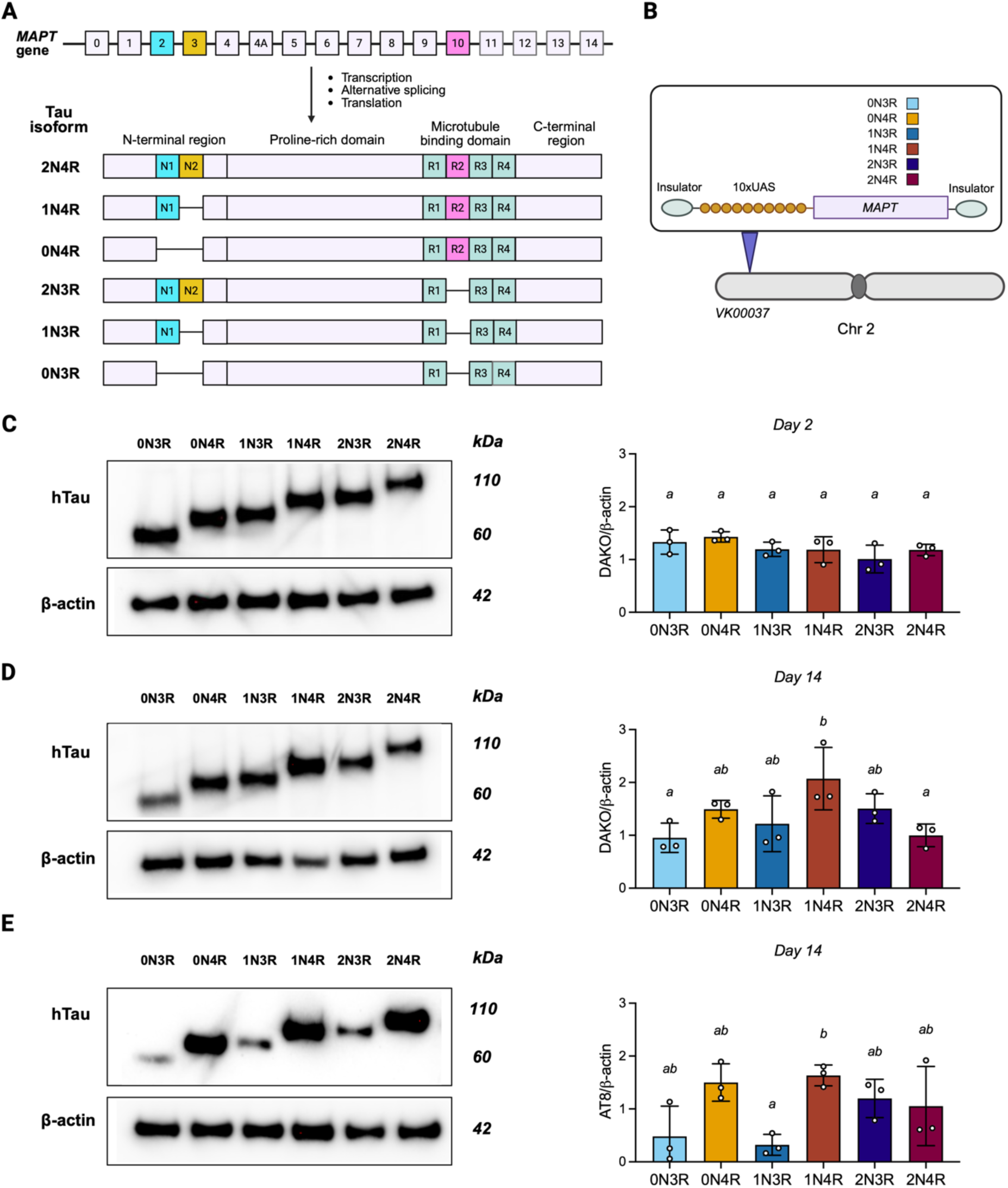
Generation and validation of transgenic *Drosophila* lines expressing the six human Tau (hTau) isoforms. **A** Schematic representation of the human *MAPT* locus and the six Tau isoforms arising from alternative splicing. **B** Overview of the six hTau constructs, each inserted at the VK00037 landing site on chromosome 2. **C-E** Pan-neuronal hTau expression was driven by elav-GAL4, with adult-restricted induction using the TARGET system (tubGal80^ts^); representative western blots are shown on the left and corresponding quantification on the right. **C** Total hTau (DAKO) levels were determined by normalisation to ß-actin levels (DAKO/ß-actin) at day 2 post-eclosion (n = 3 independent experiments, mean ± SD). Letters denote statistical groupings (one-way ANOVA with post-hoc Tukey-corrected multiple comparisons); groups sharing a letter are not significantly different (p < 0.05). **D** Total hTau at day 14 post-eclosion, showing higher levels of 1N4R accumulation compared to 0N3R and 2N4R. **E** AT8-positive phosphorylated hTau (pTau), showing higher levels for 1N4R than 1N3R.

Quantification of pan-neuronally expressed hTau by western blotting showed that the levels were comparable across isoforms at day 2 post-eclosion (Fig. 1C). However, by day 14, isoform-dependent differences in hTau accumulation were evident, with 1N4R showing higher hTau than 0N3R (p = 0.032) and 2N4R (p = 0.041) (Fig 1D). Next, we quantified the levels of phosphorylation at the AT8 epitope site (Ser202/Thr205), which has been widely used as a marker of tau pathology [18]. At day 14, AT8-positive hTau (pTau) levels differed across isoforms, with 1N4R showing higher levels than 1N3R (p = 0.036) (Fig 1E). Overall, isoform expression levels in our new hTau toolset were broadly comparable in young flies, with isoform-dependent differences in total and pTau evident at later time points.

### hTau isoforms differentially affect fly survival and behaviour

The effects of hTau on *Drosophila* lifespan have been well-documented [12], [14]. We next asked whether the pan-neuronal expression of individual hTau isoforms differentially affected lifespan. As measured by the Hazard Ratio (HR), adult expression of most hTau isoforms significantly reduced *Drosophila* lifespan compared to controls expressing ß-galactosidase (LacZ) (likelihood ratio test: χ² = 307.1, p < 0.001; Fig. 2A). Of the six isoforms, 0N4R showed the greatest toxicity (HR = 8.37, p < 0.001), followed by 2N4R (HR = 5.07, p < 0.001), 1N4R (HR = 4.18, p < 0.001), and 2N3R (HR = 2.28, p < 0.001). Expression of 0N3R and 1N3R did not reduce lifespan (see Suppl. Table 2 for all comparisons).

**Figure 2.**
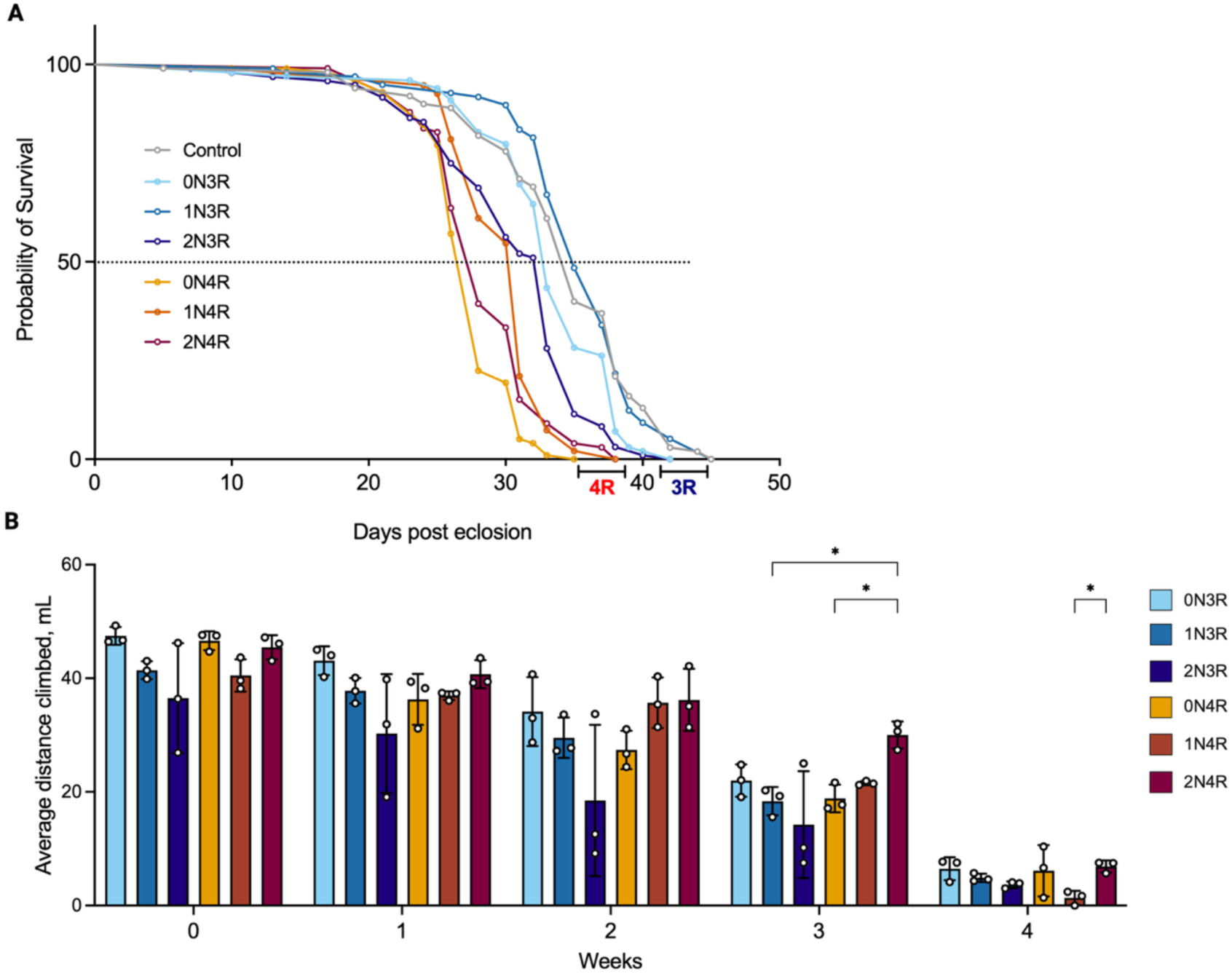
Behavioural assays demonstrate differential toxicity of the six hTau isoforms. **A** Lifespan of flies expressing each hTau isoform pan-neuronally (adult-onset induction), shown as Kaplan-Meier survival curves (n = 100 flies per genotype). Survival differences were analysed using a Cox proportional hazards model with Tukey-corrected post-hoc comparisons. 0N4R was the most toxic isoform in the lifespan assay, followed by 2N4R and 1N4R; 2N3R also reduced lifespan, whereas 0N3R and 1N3R did not. **B** Negative geotaxis (climbing) following adult-onset hTau expression in motor neurons (OK371-GAL4) (n = 3 cohorts of 10 flies per genotype). Climbing distance after 10 s was analysed by repeated-measures two-way ANOVA with Tukey-corrected post-hocs. Climbing performance was broadly comparable across isoforms at early time points. At later timepoints, flies expressing 2N4R isoform appeared to retain higher climbing performance than 0N4R and 1N3R (week 3), and 1N4R (week 4) (* p < 0.05).

Although lifespan analysis provides a measure of overall tau toxicity, it does not capture neuronal dysfunction. Behavioural impairments, including locomotor deficits, are established readouts of tau-induced neuronal dysfunction in *Drosophila* [19]. To determine whether different hTau isoforms produce different functional impairments, we expressed the isoforms in motor neurons using the *OK371*-GAL4 driver with developmental expression restricted using the Gal80^ts^ repressor and quantified climbing ability across five weeks (Fig. 2B). Climbing performance progressively worsened over time, and by week 4 flies were almost immobile. The time course of the impairment differed across isoforms (F (14.16, 33.05) = 2.45, p = 0.017), although isoform-specific differences were only evident at later time points, with 2N4R showing milder toxicity than 0N4R and 1N3R at week 3, and than 1N4R at week 4 (Fig. 2B).

### Differences in the effects of hTau isoform on degenerative phenotypes

Having established isoform-dependent effects on survival and climbing performance, we then asked how these functional outcomes relate to toxicity phenotypes elsewhere in the fly. As an initial, higher-throughput readout of tissue damage, we leveraged the adult *Drosophila* wing, which provides a rapid and quantitative *in vivo* platform for assessing tau toxicity [20]. Using engrailed-GAL4, each hTau isoform was expressed in the posterior wing compartment, with the anterior compartment serving as an internal control (Fig. 3A). As previously described in [20], Tau toxicity was quantified as the Posterior/Anterior (P/A) wing area ratio, which decreases with tissue loss due to degeneration of the targeted posterior compartment tissue. In this assay, hTau toxicity differed across isoforms (Fig. 3B; H(6) = 126.4, p < 0.0001). Damage was most pronounced with 0N4R and 1N4R, which were equally toxic (Suppl. Table 4). 0N3R and 1N3R produced more modest but still detectable reduction in wing size, while the 2N isoforms (2N3R and 2N4R) showed little impact, the latter consistent with our previous observations [20].

**Figure 3.**
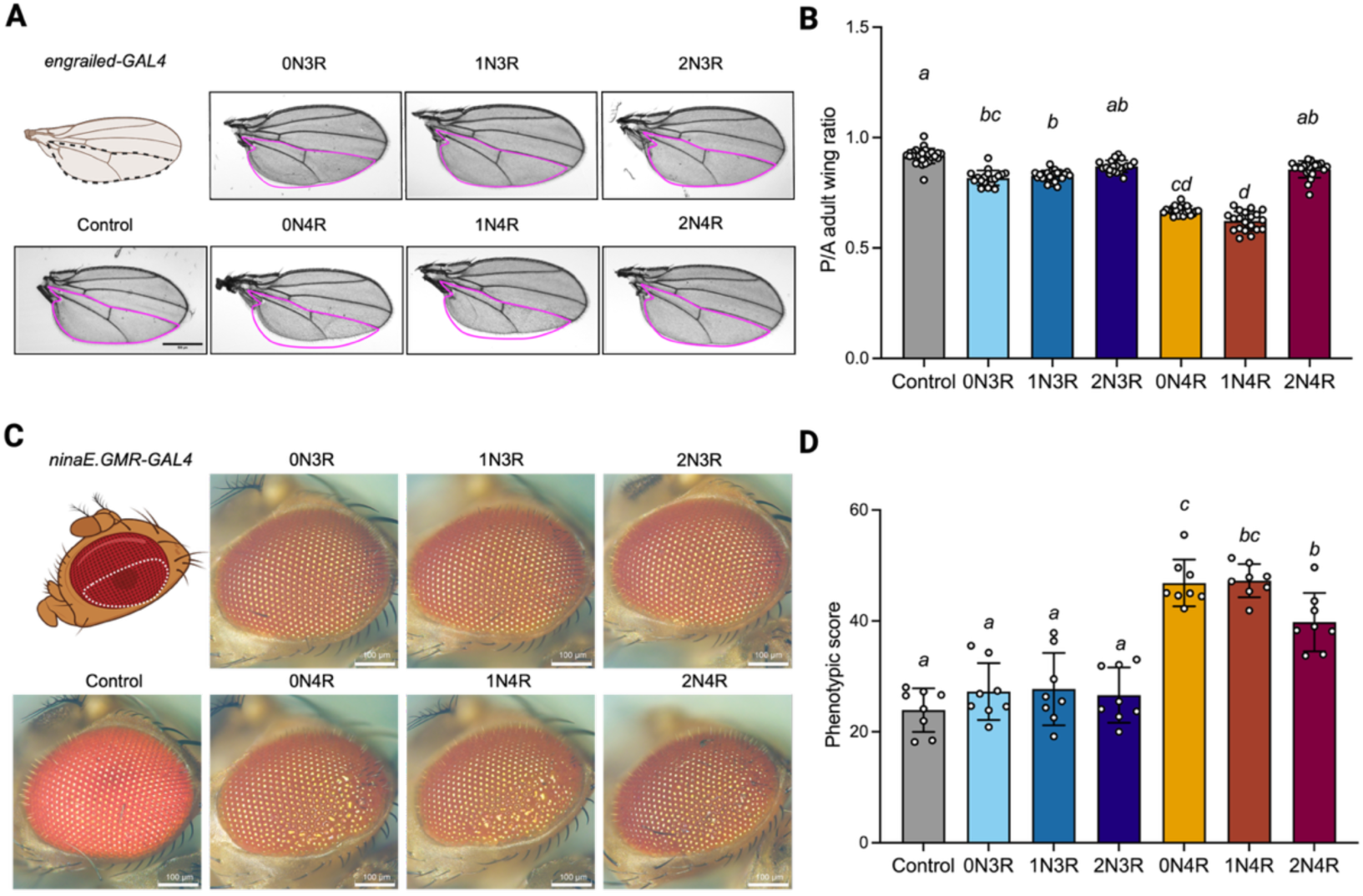
hTau isoforms cause differential patterns of toxicity depending on tissue. **A** Toxicity in the wing disc assayed by driving hTau isoform expression in the posterior compartment of the *Drosophila* adult wing using engrailed-GAL4. Flies expressing ß-galactosidase were used as controls. **B** Severity of degeneration was quantified with the posterior to anterior (P/A) compartment ratio, with lower values indicating greater impact on the posterior compartment (n = 27, 18, 23, 20, 23, 20, 21, mean ± SD). Letters denote statistical groupings (Kruskal-Wallis with Dunn’s multiple comparisons); groups sharing a letter are not significantly different. All isoforms except 2N3R and 2N4R showed some toxicity compared to ß-galactosidase (LacZ) controls. **C** Degeneration in the fly eye following hTau isoform expression driven by ninaE.GMR-GAL4, with ß-galactosidase (LacZ) controls. Images were acquired less than 24 hours post-eclosion, at 560x total magnification. **D** Degeneration was quantified using the Flynotyper ImageJ plugin and presented as a Phenotypic Score with higher values representing greater degeneration (n = 8, mean ± SD). Toxicity was observed only with 4R isoforms, with 2N4R resulting in a milder phenotype than 0N4R and 1N4R (one-way ANOVA with Tukey-corrected post-hoc tests).

To assess whether the isoform toxicity patterns observed in the wing generalise to neuronal tissue, we expressed each hTau isoform in the adult visual system using ninaE.GMR-GAL4, which drives expression in photoreceptor cells posterior to the advancing morphogenetic furrow [21] (Fig. 3C). The adult compound eye is composed of a highly ordered ommatidial lattice, making disruptions to ommatidial organisation a sensitive structural readout of degeneration [22]. In the eye, only the 4R isoforms induced ommatidial degeneration (Fig. 3D; F(6, 49) = 35.10, p < 0.0001), whereas 3R isoforms did not produce detectable damage (Suppl. Table 5). Among the 4R isoforms, 2N4R showed milder toxicity than the others.

### Tau isoforms and selective vulnerability

In Sivanantharajah *et al.* (2025) [16], we expressed hTau0N3R in small homogeneous populations (hemilineages) of neurons in the adult *Drosophila* CNS, and found that some neurons were highly vulnerable to this isoform, leading to severe degeneration, whilst others remained unaffected. To determine if selective vulnerability is driven by the unique properties of individual hTau isoforms or whether all isoforms induce comparable pathology within the same neuron type, we expressed the six hTau isoforms in neurons previously determined to be resilient (19B) and vulnerable (6A) to 0N3R [23]. To assess effects of ageing, we compared young adults shortly after eclosion (day 0) with individuals aged up for three weeks (day 21).

The 19B hemilineage comprises a small cholinergic population of interneurons originating from the neuroblast (NB) 6-2 [24]. Given the previously demonstrated resilience of 19B to the 0N3R isoform [16], we assessed whether this effect extended to all six hTau isoforms. To analyse the effects of isoforms on overall 19B neuronal morphology, we labelled cell membranes using membrane-associated green fluorescent protein (mCD8::GFP) and quantified total neuronal area as a morphological readout for degeneration (Fig. 4A). Shortly after eclosion (day 0), morphology of resilient 19B neurons was largely unaffected: none of the hTau isoform-expressing groups showed a significant reduction in neuronal area relative to control (Fig. 4B). At this early timepoint, we observed small differences in neuronal area between specific isoform pairs (0N3R vs 2N3R, 0N4R vs 2N3R, 1N4R vs 2N3R; Suppl. Table 6). In contrast, by day 21 post-eclosion, all hTau isoforms produced a comparable reduction in neuronal area relative to the control (p < 0.003). Thus, 19B neurons show early resilience to hTau-mediated degeneration, but ageing shifts this lineage towards a common degenerative outcome regardless of isoform. This contrasts with prior work reporting largely normal 19B morphology at 3 weeks following hTau0N3R expression [16]; however, this may be accounted for by the use of a completely different transgene with differing expression levels.

**Figure 4.**
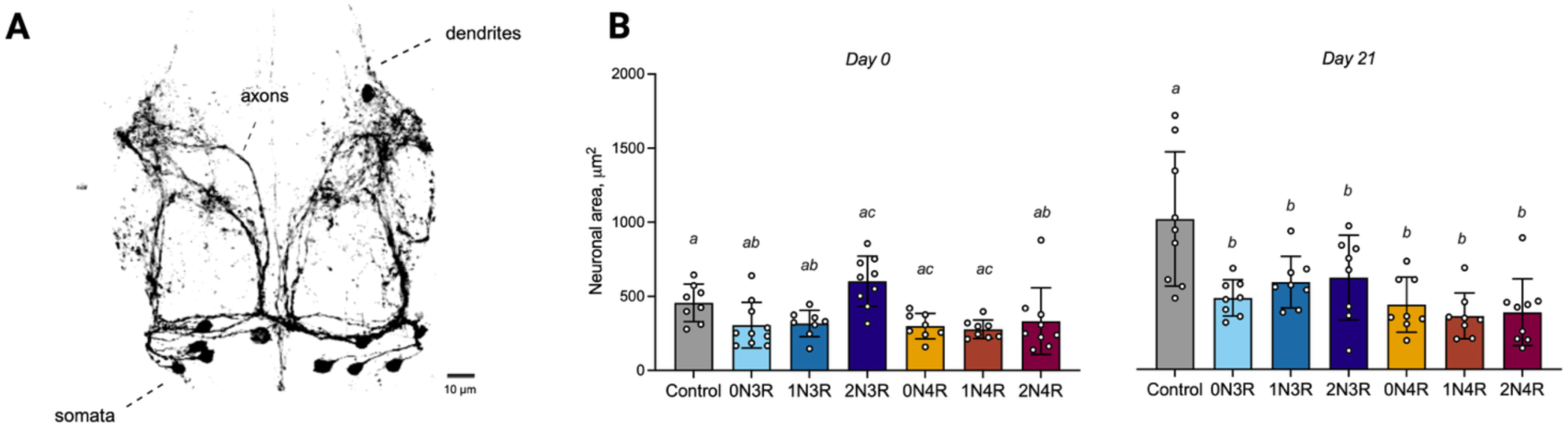
hTau expression ultimately leads to degeneration even in resilient neurons. **A** Morphology of neurons previously observed to show resilience to the 0N3R isoform (19B) visualised using mCD8::GFP. **B** Quantification of neuronal area at two timepoints, day 0 and 21 post-eclosion (n = 8-10, mean ± SD). ß-galactosidase (LacZ) expression was used as a control. Letters denote statistical groupings; groups sharing a letter are not significantly different (two-way ANOVA with Tukey-corrected post-hoc tests, p < 0.05). hTau expression only resulted in degeneration at day 21, with all isoforms showing similar toxicity.

To explore whether the resilience-to-neurodegeneration transition is accompanied by changes in tau burden [25], we quantified total hTau and AT8 phosphorylation (pTau) levels in 19B neurons. Total hTau levels showed no age-dependent increase and were broadly comparable across isoforms and time points (Suppl. Fig. 1A), suggesting that the loss of 19B resilience with age is not readily explained by isoform-dependent differences in hTau accumulation. In contrast, pTau levels were similar across isoforms shortly after eclosion but differed later in life. At day 21, two 4R isoforms (0N4R and 1N4R) had higher pTau levels than the corresponding 3R isoforms (0N3R and 1N3R) (Suppl. Fig. 1B). Together, this indicates that, in the 19B hemilineage, total hTau levels are comparable across isoforms, whereas AT8 phosphorylation differences emerge over time without correlating with degeneration severity.

We next examined the vulnerable 6A hemilineage to determine whether hTau isoforms differentially affect degeneration severity in a susceptible neuron type. The 6A hemilineage comprises GABAergic interneurons involved in processing sensory feedback during flight-related motor control (Fig. 5A). We measured total neuronal area of 6A neurons as a readout for morphological degeneration (Fig. 5B). In both young and old adults, expression of all hTau isoforms except 2N3R resulted in significant reduction in neuronal area relative to control (p ≤ 0.0390; Suppl. Table 7), indicative of gross neuronal degeneration. Beyond this, isoform-specific differences were limited, with 2N4R showing higher toxicity than 2N3R (p = 0.033; Suppl. Table 7). Together, these data indicate that 6A neurons are broadly susceptible to tau-mediated degeneration across most isoforms, with significant reductions in neuronal area apparent since shortly after eclosion. To assess whether hTau abundance or phosphorylation state relates to degeneration severity in 6A neurons, we quantified total hTau and pTau levels (Suppl. Fig. 1C,D) and found that levels of both increased over time but remained broadly comparable across isoforms. Therefore, what small isoform-dependent differences that exist in causing degeneration in 6A neurons, cannot be completely explained by levels of hTau or pTau.

**Figure 5.**
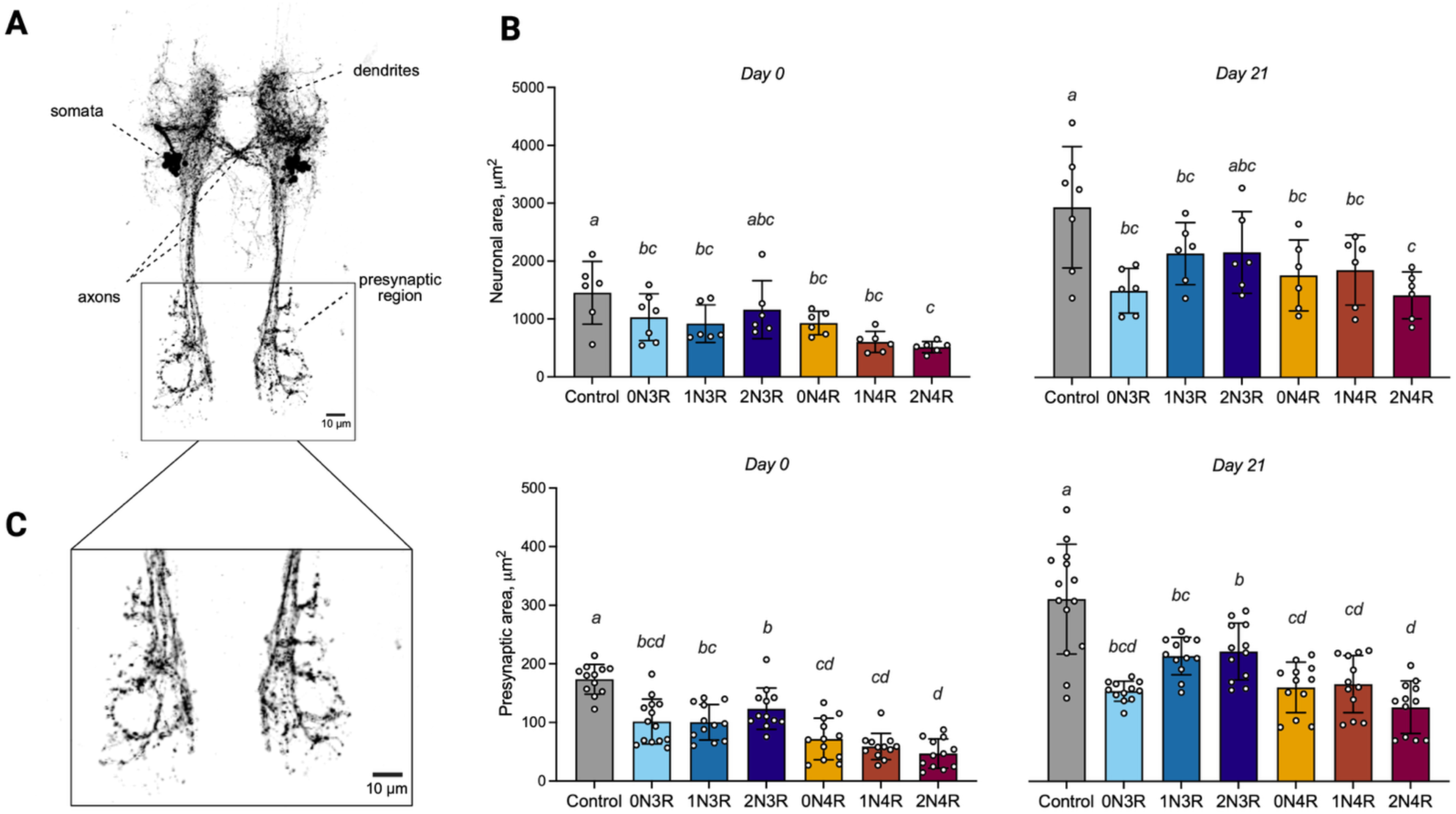
Isoform-specific patterns of hTau toxicity in vulnerable neurons. **A** Neuronal morphology of the vulnerable 6A hemilineage visualised using mCD8::GFP. **B** Neuronal area was quantified at day 0 and day 21 post-eclosion (n = 6-7, mean ± SD). Letters denote statistical groupings; groups sharing a letter are not significantly different (two-way ANOVA with post-hoc Tukey-corrected multiple comparisons; p < 0.05). Degeneration was observed at both time points for all isoforms except 2N3R. **C** Presynaptic terminal area of 6A neurons was measured as a subcellular readout of toxicity. The presynaptic region used in the analysis is highlighted on the left. Although all hTau isoforms resulted in degeneration, 4R isoforms exhibited greater toxicity.

Since the 6A neurons are highly polarised with long, regionally specialised structures, whole-cell morphology provides only a coarse readout of degeneration. In our previous work, one of the prominent pathological features in 6A neurons was a reduction in size of distal presynaptic terminals, indicating sub-cellular, regional sensitivity to degeneration. This is consistent with the broader literature showing that synaptic dysfunction often precedes major structural loss and can serve as a more sensitive indicator of tau-mediated pathology [26]. We quantified presynaptic area in 6A neurons (Fig. 5C) and found that all hTau isoforms resulted in reduced presynaptic area relative to control (p ≤ 0.002; Suppl. Table 8). We also observed isoform-specific differences in toxicity: 2N4R produced the greatest presynaptic loss, followed by 0N4R and 1N4R. 0N3R showed an intermediate phenotype, while 1N3R and 2N3R were least toxic. Notably, synaptic toxicity was detectable even for 2N3R, which showed little impact on gross morphology, underscoring the sensitivity of this presynaptic readout. Overall, our analysis of distinct neuronal subpopulations showed that hTau accumulation and AT8-positive phosphorylation did not consistently predict the extent of degeneration, indicating that neuronal responses to hTau are shaped by other cell-intrinsic factors.

## Discussion

Despite tau pathology being a hallmark of many neurodegenerative diseases, the functional differences between tau isoforms and their relative contributions to disease progression remain incompletely understood. This knowledge gap is especially relevant given that distinct tauopathies are characterised by different isoform compositions, with some showing predominant accumulation of 3R tau, 4R tau, or a mixture of both, suggesting that isoform balance may shape clinicopathological heterogeneity. However, most experimental models, including *Drosophila* tauopathy models, typically express a single hTau isoform, and isoform choice varies widely between studies, often without a clear rationale [27]. This reductionist approach limits direct, within-systems comparisons and may therefore not capture the full spectrum of isoform-specific toxicity observed in human disease. A comprehensive study of isoform-specific effects is ultimately essential for understanding disease heterogeneity and designing targeted interventions. Our study provides a systematic analysis of the toxicity of all six hTau isoforms using a newly developed set of *Drosophila* lines that, by restricting genetic variability, allow clear delineation of isoform-specific differences (summarized in Fig. 6).

**Figure 6.**
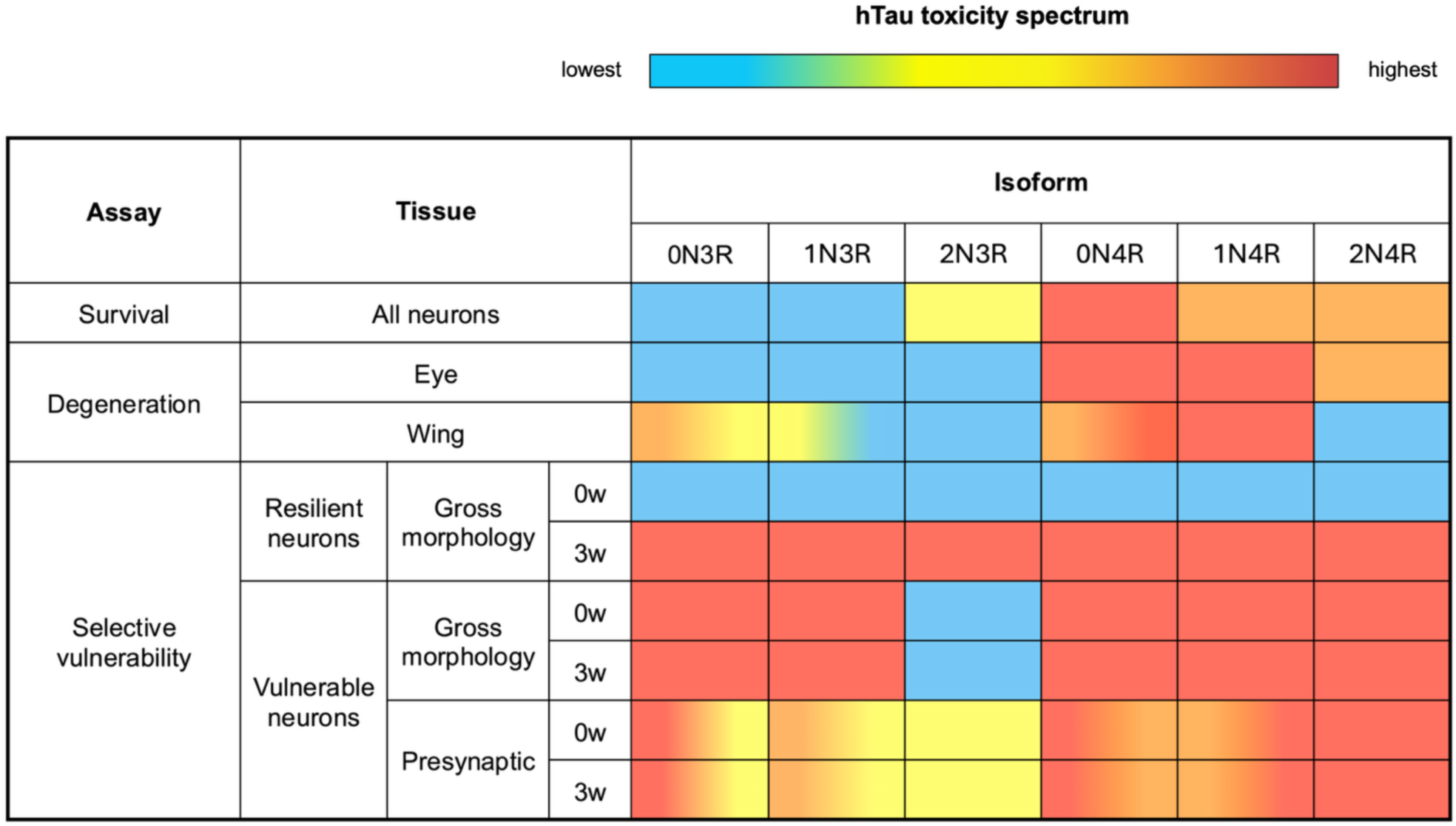
Summary of results: context-dependent toxicity of the hTau isoforms. Each cell is coloured according to the post-hoc statistical groupings within that assay (p < 0.05), with colours chosen to reflect relative toxicity based on the assay metric (e.g., reduced lifespan, increased degeneration, higher phenotypic score, decreased neuronal area). Red indicates the most toxic group, and blue denotes no significant difference from the ß-galactosidase (LacZ) control. Where a cell contains two colours, the isoform toxicity is not significantly different from either colour group, although these groups differ from each other. Note, no continuous scoring system was used and no numerical values were assigned to the colours.

We show that, overall, 4R hTau isoforms exhibit markedly higher toxicity than the 3R isoforms. Lifespan analysis revealed a significant reduction with the three 4R isoforms and only 2N3R (Fig. 2A). Interestingly, this contrasts with findings from Fernius *et al.* (2017), who observed broadly similar effects of all hTau isoforms on lifespan, except for elevated toxicity of 2N3R. Our toxicity assays using the eye and wing readouts also pointed to generally higher toxicity of 4R isoforms. Previous work has shown that expression of hTau during development can be important for triggering degenerative phenotypes [13]. Consistent with this, developmental expression of 4R isoforms driven by GMR-GAL4 resulted in severe degeneration whereas 3R expression did not elicit defects (Fig. 3C). Since the only hTau isoform expressed during human brain development is 0N3R, the extra microtubule-binding repeat in the 4R isoforms may interact with cytoskeletal proteins, drastically altering normal development [28]. Nevertheless, the reduction in lifespan we observed when 4R tau expression was restricted to adults using the TARGET system, indicates that developmental expression is not essential for tau-mediated toxicity and can occur in adult flies as a function of age.

The higher toxicity of 4R isoforms aligns well with the isoform imbalance observed in human tauopathies. Primary tauopathies, such as PSP, CBD, and many others are characterised by the pathological accumulation of 4R tau, whereas 3R-dominant pathology is rare [29, 30]. A possible structural feature that could contribute to this 3R-4R differential toxicity is the presence of an additional hexapeptide aggregation motif, VQIINK, in the R2 repeat [31]. VQIINK can form highly stable steric zippers leading to strong self-aggregation capacity [32]. Although VQIVYK is essential for filament formation, VQIINK deletion has been linked to reduced tau assembly, implicating VQIINK as a potential driver of the enhanced 4R toxicity [33].

However, the pattern of toxicity across our phenotypic measures was more nuanced than a simple 3R-4R dichotomy, with meaningful isoform-specific differences emerging across assays. Analysis of hTau-induced functional deficits using negative geotaxis revealed some isoform-specific effects, although these differences only emerged at later weeks (Fig. 2B). Notably, the 2N4R isoform had relatively preserved climbing performance at later time points despite showing reduced survival, suggesting that locomotor dysfunction and organismal viability do not necessarily deteriorate in parallel. We also did not detect increased toxicity of 0N3R relative to 0N4R, in contrast to findings reported by Sealey *et al.* (2017), indicating that isoform-dependent effects can vary by phenotype and experimental context [12].

Importantly, our degeneration assays in the wing and the eye showed considerable variation in toxicity within both the 3R and 4R groups. In both tissues, 2N isoforms containing both N-terminal inserts (2N) sometimes produced no obvious degeneration (2N3R in the eye, and 2N3R/2N4R in the wing). This suggests important modulatory contributions of the regions outside the microtubule-binding domains. The functional profile of the N-terminal region appears highly complex, with documented involvement in modulating microtubule stabilisation (e.g., bundle formation/spacing), regulation of axonal transport, and interactions with plasma membrane [34], [35]. Structurally, the N-terminal region is intrinsically disordered, which may facilitate diverse intermolecular interactions, influencing the ability of tau to engage in complex regulatory networks or dynamic assemblies. Tau structural models, such as the ‘paperclip’ conformation, propose that the N- and the C-termini fold back onto the repeat domain, with the number of the N-terminal inserts modulating the protein conformation [36], [37]. Together, these observations support the idea that the N-terminal composition can tune the conformational ensemble of tau and its interaction landscape, shaping whether toxicity leads to overt degeneration or is buffered despite belonging to the same 3R or 4R group.

Taken together, our findings also suggest that isoform toxicity is shaped by tissue-specific factors. In the wing, damage was most pronounced with the 0N4R and 1N4R isoforms, whereas 0N3R and 1N3R were less toxic but still produced a detectable posterior-compartment phenotype (Fig. 3B). In contrast, in the neuronal context in the eye, only 4R isoforms induced ommatidial degeneration, while 3R isoforms produced no detectable damage (Fig. 3D). Overall, these tissue-specific differences in toxicity indicate that local cellular environments may influence the effects of tau. One possible explanation is that tissues differ in the relative abundance and activity of tau modifiers and interactors (e.g., phosphorylation machinery, cytoskeletal pathways, quality-control pathways etc.). These tissue-specific factors are likely to shape selective vulnerability: the same tau isoform may be tolerated in some cell populations yet drive degeneration in others. Isoform-dependent toxicity would then emerge from the interaction between tau structure and cell-type-specific molecular landscapes.

To study neurodegeneration in the context of selective vulnerability, we restricted hTau isoform expression to small, highly-homogeneous neuronal populations in the fly CNS that were previously identified as vulnerable or resilient to the 0N3R hTau isoform [16]. This approach offers several advantages. First, these hemilineages are anatomically well-characterised, enabling reliable identification of distinct sub-cellular compartments. Another major advantage of our system is its resolution. Given that degeneration may be first detected in particularly vulnerable neuronal regions (e.g., synaptic terminals), subcellular analysis of neuronal morphology may reveal differences that are not captured by gross area measurements. The strengths of this system were further enhanced by our hTau isoform panel, in which each transgene was inserted at a defined landing site to reduce expression-related confounds and improve comparability between the lines.

The resilient 19B and vulnerable 6A hemilineages showed fundamentally different temporal responses to hTau expression. In 19B neurons, isoform-specific toxicity was minimal early after eclosion: no isoform produced clear degenerative effects shortly after eclosion, and only subtle differences were detected between a small number of isoform pairs (Fig. 5). By three weeks, however, this early resilience was lost and all isoforms converged on a similar toxic outcome, with little evidence for sustained isoform selectivity. This suggests that in 19B neurons, intrinsic factors may initially constrain toxicity, but this protection diminishes with age, implying that resilience is transient and breaks down as tau burden increases over time (i.e., with age). This transition from resistance to vulnerability may parallel the hierarchical progression described by Braak staging [25], where tau pathology emerges in selective regions before spreading to broader cortical territories. Age-dependent breakdown of intrinsic constraints could therefore help explain why certain neuronal populations become permissive to pathology only at later stages of disease.

In contrast, 6A neurons showed a rapid and severe response to hTau expression, with degeneration evident from the earliest time point and persisting at three weeks (Fig. 6). Isoform-dependent differences were limited, as all isoforms were toxic except for 2N3R, and 2N4R showed greater toxicity than 2N3R. Importantly, this pattern could not be explained by differences in total hTau abundance or AT8-phosphorylated tau levels alone. Rather than overall tau load or phosphorylation at a single epitope, toxicity likely depends on how tau behaves within a given neuronal context - its subcellular localisation, conformational state, broader post-translational modification profile, and interactions with cell-type-specific pathways.

This underscores the value of subcellular analysis, as context-dependent differences in tau behaviour may become more apparent in particularly vulnerable neuronal regions. In the presynaptic compartment, we observed clear isoform-dependent stratification in toxicity. 2N4R was the most toxic isoform, followed by modest toxicity of 0N4R and 1N4R, intermediate toxicity of 0N3R, and mild toxicity of 1N3R and 2N3R (Fig. 5C). These findings directly further our previous work on selective vulnerability and highlight that both neuronal identity and isoform specificity determine the severity of pathological response. Furthermore, the sensitivity of 6A presynaptic structures to tau-mediated degeneration underscores the value of subcellular resolution in investigating early neuronal pathology.

The hTau isoform line set generated here offers a tractable *in vivo* platform for dissecting isoform-specific tau toxicity. Using this platform, we identify a clear context-dependent split in isoform toxicity, with 4R isoforms generally conferring higher toxicity than 3R isoforms, yet with substantial heterogeneity across measures, tissues, and neuronal populations. Future work could focus on investigating the 4R-specific features (e.g., R2 including the VQIINK motif) and systematic analysis of isoform-specific post-translational modification landscapes that may modulate subcellular localisation, aggregation rate, and neurotoxicity. More broadly, this set of six hTau isoform *Drosophila* lines provide a foundation for scalable candidate and unbiased modifier approaches to map the cellular pathways that buffer or exacerbate isoform-specific toxicity.

## Materials and methods

### Drosophila culture

Flies were raised on a standard diet of cornmeal with a 12h light: 12h dark cycle. For experiments using the TARGET system [38] to restrict gene expression to adult neurons (e.g., western blotting, survival, negative geotaxis), crosses were reared at 18°C and flies of relevant genotypes moved to 29°C upon eclosion. Crosses using the ninaE.GMR-GAL4 driver to examine rough eye phenotypes were reared at 25°C. Details of all *Drosophila* stocks used in this study are provided in Suppl. Table 1.

### Transgenic hTau isoform flies

To create new hTau expressing lines, DNA sequences for each of the six hTau isoforms were placed downstream of ten GAL4 upstream activation sequences (10xUAS). These constructs were insulated with two *gypsy* insulators and inserted into chromosome 2 at the same landing site, VK00037 (Fig. 1B). Cloning was completed by GenScript (NJ, USA) and the injection and creation of stable transgenic lines by GenomeProlab (Montreal, Canada).

### Western blotting to quantify transgene expression

To test if new transgenic lines expressed hTau at similar levels, expression of each isoform was driven in using the pan-neuronal driver, *elav-GAL4* and *tubGal80^ts^* to restrict expression to adult flies. Crosses were set up at 18°C and upon eclosion, male *Drosophila* were collected and aged at 29°C for two days. Flies were decapitated and stored at –80°C until use. Frozen heads were placed in ice-cold homogenisation buffer (HB): 10mM Tris, 150mM NaCl, 5mM EDTA, 20% v/v glycerol, protease and phosphatase inhibitors (1:100). Fly heads were manually homogenised for 3 minutes in HB (5 uL/head). Lysates were centrifuged at 10000 rpm at 4°C for 5 minutes. The supernatant was removed, mixed with NuPAGE LDS sample buffer (NP0007, Thermofisher) and DTT and boiled at 95°C for 5 minutes. Western blotting was performed using standard protocols [39].

Primary antibodies used in this study were: rabbit anti-hTau (Agilent – DAKO, Cat. no. A002401-2) at 1:15,000 dilution, mouse anti-phospho-Tau (Ser202, Thr205) antibody (AT8, Fisher, Cat. no. 10599853) at 1:1000 dilution, and mouse anti-β-actin antibody (AbCam, ab8224) at 1:3000 dilution. Secondary antibodies used were: anti-rabbit HRP-conjugated (Invitrogen, 31460) and anti-mouse HRP-conjugated (Invitrogen, 31430). Antibody-antigen complexes were detected using Chemiluminescent Peroxidase Substrate-3 (Sigma-Aldrich, CPS350-1KT). Blots were imaged on a Bio-Rad ChemiDoc system. Protein expression levels were quantified by densitometric analysis of bands using ImageJ (i.e., FIJI, (http://rsbweb.nih.gov/ij/)).

### Survival assay

To determine the effect of each isoform on fly lifespan, the TARGET system was used with pan-neuronal elav-GAL4 to restrict expression of transgenes to adult flies [17]. elav-GAL4, tubGal80^ts^ flies were crossed to each of the six UAS-hTau isoform lines and UAS-lacZ, used as a protein overexpression control. Crosses were maintained at 18°C to suppress any transgene expression. Upon eclosion, male progeny was collected and transferred to 29°C (10 flies per vial). Flies were transferred to new vials every 2-3 days and the number of surviving flies was recorded. Kaplan-Meier curves were generated using GraphPad Prism (version 10.6; GraphPad Software, 2025). Survival was analysed in R using a Cox proportional hazards model: Surv(Day, Status)∼Genotype+frailty(Vial).

### Negative geotaxis assay

To examine the effects of isoform expression on climbing behaviour, transgenic *UAS-hTau Drosophila* lines were crossed to the motor neuron-specific, OK371-GAL4 driver. Expression of hTau was restricted to adults using the TARGET system [17]. Male progeny were collected and transferred to 29°C (10 flies per vial). On the day of the experiment, flies were flipped into cylinders and allowed to acclimatise to the room temperature for one hour. The vials were gently tapped, and the distance climbed by each fly after 10 seconds was recorded. This procedure was repeated three times with 30 second breaks in between the measurements.

### Drosophila wing assay

To examine hTau isoform toxicity in a second tissue, we drove expression of isoforms using the engrailed-GAL4 driver [40], which drives transgene expression in the posterior compartment of the wing. Flies were reared at 25°C, and male progeny was collected up to 48 hours after eclosion and frozen. Wings were removed and imaged using a Leica Stereoscopic microscope (LAS Software). The ratio of posterior to anterior wing compartments was calculated as described in [20] with Fiji.

### Eye phenotype analysis

To determine if different hTau isoforms could elicit a rough eye phenotype, we used the ninaE.GMR-GAL4 driver. *Drosophila* eyes were imaged using a DSX-1000 microscope (Olympus) with a 40x objective and the final magnification of 560x. Flynotyper plugin in ImageJ (https://flynotyper.sourceforge.net/imageJ.html) was used to calculate phenotypic scores, with higher values indicating greater degeneration.

### VNC tissue immunohistochemistry and confocal microscopy

Flies for these experiments were set at 18°C, and the progeny was transferred to 29°C after eclosion. Ventral nervous systems (VNSs) were dissected and processed as described in [24]. Primary antibodies used were: 1:1000 chicken anti-GFP (Invitrogen-Molecular Probes, Cat. no. A10262), 1:15000 rabbit anti-hTau (Agilent-DAKO, Cat. no. A002401-2), 1:125 mouse anti-lacZ (J1E7, DSHB); 1:1000 mouse anti-phospho-Tau (Ser202, Thr205) antibody (AT8, Fisher, Cat. no. 10599853). Relevant secondary antibodies were used at 1:500. Confocal microscopy images were taken using a Zeiss LSM 710 confocal microscope. Images used for signal quantification were obtained at 40x oil magnification using identical image acquisition parameters.

### Analysis of neuronal morphology

Neuronal morphology was visualised using the mCD8-GFP signal. Neuronal area was measured using the maximum intensity projections of the neuronal images. For the subcellular morphological analysis, two measurements per VNS were made, with six VNSs analysed per isoform. Statistical analysis was performed in R using a linear mixed effects model (package ‘lme4’, v.1.1.35.5). Isoform, time, and their interaction were modelled as fixed effects, and the ID of the VNS analysed was included as a random effect: lmer(Area ∼ Isoform*Time + (1|Subject)). hTau and p-Tau levels were measured as corrected total cell fluorescence (CTCF), where CTCF = Integrated Density – (Area of selected cell x Mean fluorescence of background readings) [41].

### Statistical analysis

Unless stated otherwise, all statistical analyses were performed in GraphPad Prism (version 10.6; GraphPad Software, 2025). A Kruskal-Wallis test with post-hoc Dunn’s tests were used to evaluate the differences in P/A wing ratio (Suppl. Table 5). Repeated measures two-way ANOVA with Tukey-corrected post-hoc tests was used to analyse the effects of isoforms on climbing ability (Suppl. Table 3). One-way ANOVAs with Tukey-corrected post-hoc tests were used to assess the differences in the eye phenotypic score between the isoforms (Suppl. Table 4). Two-way ANOVAs with Tukey-corrected post-hoc tests were used to analyse the effects of isoform expression, time, and their interaction on (i) neuronal area, (ii) hTau, and (iii) p-tau levels (Suppl. Fig. 1).

## Supporting information

All supplementary materials

