## Supplementary material for "Context-dependent toxicity of human Tau isoforms in a *Drosophila* tauopathy model": All supplementary materials

**Supplementary table 1.** *Drosophila* stocks used in this study.

| Fly line | Genotype | Source | Use |
|---|---|---|---|
| hTau0N3R.VK37 | +*; UAS-hTau0N3R.VK00037/CyO;*+ | This paper | All assays |
| hTau0N4R.VK37 | +*;UAS-hTau0N4R.VK00037/CyO;*+ | This paper | All assays |
| hTau1N3R.VK37 | +*; UAS-hTau1N3R.VK00037/CyO* | This paper | All assays |
| hTau1N4R.VK37 | *; UAS-hTau1N4R.VK00037/CyO* | This paper | All assays |
| hTau2N3R.VK37 | *; UAS-hTau2N3R.VK00037/CyO* | This paper | All assays |
| hTau2N4R.VK37 | *; UAS-hTau2N4R.VK00037/CyO* | This paper | All assays |
| lacZ | *; UAS-lacZ/CyO; MKRS/TM6B* | BDSC no. 8529 | Control |
| elav-GAL4, tubGal80$^{ts}$ | *elav-GAL4; + ;tubGal80*$^{ts}$ | BDSC no. 458 | Survival, western blotting |
| OK371, tubGal80$^{ts}$ | *; OK371; tubGal80*$^{ts}$ | BDSC no. 26160 | Negative geotaxis |
| GMR-GAL4 | *; GMR-Gal4/CyO* | BDSC no. 9146 | Eye phenotype analysis |
| engrailed-GAL4 | *; engrailed2.4-GAL4e16E* | BDSC no. 30564 | Wing assay |
| 6A-GAL4.GFP | *; 6A-GAL4, mCD8::GFP/CyO; 6A-GAL4/MKRS* | Modified from 6A-GAL4 (SS26967, FlyLight, Janelia) | Selective vulnerability |
| 19B-GAL4.GFP | *; 6A-GAL4, mCD8::GFP/CyO; 6A-GAL4/MKRS* | Modified from 19B-GAL4 (SS20748, FlyLight, Janelia) | Selective vulnerability |

**Supplementary table 2.** Tukey-corrected post-hoc multiple comparisons for survival assay

| | 0N3R | 1N3R | 2N3R | 0N4R | 1N4R | 2N4R |
|---|---|---|---|---|---|---|
| Control | 0.1174 | 0.9977 | <.0001 | <.0001 | <.0001 | <.0001 |
| 0N3R | | 0.0248 | 0.0424 | <.0001 | <.0001 | <.0001 |
| 1N3R | | | <.0001 | <.0001 | <.0001 | <.0001 |
| 2N3R | | | | <.0001 | 0.0008 | <.0001 |
| 0N4R | | | | | <.0001 | 0.0099 |

| | | | | | | |
|---|---|---|---|---|---|---|
| 1N4R | | | | | | 0.8395 |

**Supplementary table 3.** Tukey-corrected post-hoc multiple comparisons for negative geotaxis assay (all timepoints shown)

Week 0

| | 1N3R | 2N3R | 0N4R | 1N4R | 2N4R |
|---|---|---|---|---|---|
| 0N3R | 0.0506 | 0.5439 | 0.9786 | 0.1407 | 0.7696 |
| 1N3R | | 0.9263 | 0.0907 | 0.9926 | 0.2809 |
| 2N3R | | | 0.5953 | 0.9691 | 0.6733 |
| 0N4R | | | | 0.1927 | 0.9652 |
| 1N4R | | | | | 0.3331 |

Week 1

| | 1N3R | 2N3R | 0N4R | 1N4R | 2N4R |
|---|---|---|---|---|---|
| 0N3R | 0.2516 | 0.4995 | 0.3880 | 0.1592 | 0.6859 |
| 1N3R | | 0.8091 | 0.9901 | 0.9795 | 0.6746 |
| 2N3R | | | 0.9183 | 0.8515 | 0.6287 |
| 0N4R | | | | 0.9995 | 0.6859 |
| 1N4R | | | | | 0.3670 |

Week 2

| | 1N3R | 2N3R | 0N4R | 1N4R | 2N4R |
|---|---|---|---|---|---|
| 0N3R | 0.8454 | 0.5498 | 0.6097 | 0.9985 | 0.9964 |
| 1N3R | | 0.7423 | 0.9613 | 0.5190 | 0.5705 |
| 2N3R | | | 0.8458 | 0.4705 | 0.4591 |
| 0N4R | | | | 0.2907 | 0.3596 |
| 1N4R | | | | | >0.9999 |

Week 3

| | 1N3R | 2N3R | 0N4R | 1N4R | 2N4R |
|---|---|---|---|---|---|
| 0N3R | 0.6104 | 0.7510 | 0.7061 | 0.9994 | 0.1087 |
| 1N3R | | 0.9597 | 0.9998 | 0.4699 | 0.0239 |
| 2N3R | | | 0.9406 | 0.7621 | 0.3170 |
| 0N4R | | | | 0.5588 | 0.0268 |
| 1N4R | | | | | 0.0820 |

Week 4

| | 1N3R | 2N3R | 0N4R | 1N4R | 2N4R |
|---|---|---|---|---|---|
| 0N3R | 0.7825 | 0.4203 | >0.9999 | 0.1298 | 0.9991 |
| 1N3R | | 0.3728 | 0.9928 | 0.0956 | 0.2660 |

| | | | | | |
|---|---|---|---|---|---|
| 2N3R | | | 0.9061 | 0.2593 | 0.0800 |
| 0N4R | | | | 0.5966 | 0.9994 |
| 1N4R | | | | | 0.0251 |

**Supplementary table 4.** Dunn's multiple comparisons for the P/A adult wing ratio

| | 0N3R | 1N3R | 2N3R | 0N4R | 1N4R | 2N4R |
|---|---|---|---|---|---|---|
| Control | <0.0001 | <0.0001 | 0.5078 | <0.0001 | <0.0001 | 0.0883 |
| 0N3R | | >0.9999 | 0.1632 | 0.1237 | 0.0054 | 0.6301 |
| 1N3R | | | 0.3888 | 0.0131 | 0.0003 | >0.9999 |
| 2N3R | | | | <0.0001 | <0.0001 | >0.9999 |
| 0N4R | | | | | >0.9999 | <0.0001 |
| 1N4R | | | | | | <0.0001 |

**Supplementary table 5.** Tukey-corrected post-hoc multiple comparisons for eye degeneration

| | 0N3R | 1N3R | 2N3R | 0N4R | 1N4R | 2N4R |
|---|---|---|---|---|---|---|
| Control | 0.8070 | 0.7006 | 0.9174 | <0.0001 | <0.0001 | <0.0001 |
| 0N3R | | >0.9999 | >0.9999 | <0.0001 | <0.0001 | <0.0001 |
| 1N3R | | | 0.9993 | <0.0001 | <0.0001 | 0.0002 |
| 2N3R | | | | <0.0001 | <0.0001 | <0.0001 |
| 0N4R | | | | | >0.9999 | 0.0728 |
| 1N4R | | | | | | 0.0469 |

**Supplementary table 6.** Tukey-corrected post-hoc multiple comparisons for the neuronal area of the 19B resilient neurons

Week 0

| | 0N3R | 1N3R | 2N3R | 0N4R | 1N4R | 2N4R |
|---|---|---|---|---|---|---|
| Control | 0.7198 | 0.8250 | 0.7712 | 0.7320 | 0.6005 | 0.8824 |
| 0N3R | | >0.9999 | 0.0359 | >0.9999 | >0.9999 | >0.9999 |
| 1N3R | | | 0.0761 | >0.9999 | 0.9998 | >0.9999 |
| 2N3R | | | | 0.0485 | 0.0268 | 0.0936 |
| 0N4R | | | | | >0.9999 | 0.9999 |
| 1N4R | | | | | | 0.9980 |

Week 21

| | 0N3R | 1N3R | 2N3R | 0N4R | 1N4R | 2N4R |
|---|---|---|---|---|---|---|
| Control | <0.0001 | 0.0009 | 0.0028 | <0.0001 | <0.0001 | <0.0001 |
| 0N3R | | 0.9455 | 0.8365 | 0.9994 | 0.9022 | 0.9600 |
| 1N3R | | | >0.9999 | 0.7617 | 0.3013 | 0.4034 |
| 2N3R | | | | 0.5725 | 0.1687 | 0.2377 |
| 0N4R | | | | | 0.9900 | 0.9986 |

| | | | | | | |
|---|---|---|---|---|---|---|
| 1N4R | | | | | | >0.9999 |

**Supplementary table 7.** Tukey-corrected post-hoc multiple comparisons for the neuronal area of the 6A vulnerable neurons

| | 0N3R | 1N3R | 2N3R | 0N4R | 1N4R | 2N4R |
|---|---|---|---|---|---|---|
| Control | 0.0006 | 0.0390 | 0.1702 | 0.0029 | 0.0004 | <0.0001 |
| 0N3R | | 0.8754 | 0.5155 | 0.9997 | >0.9999 | 0.8036 |
| 1N3R | | | 0.9965 | 0.9796 | 0.8107 | 0.1453 |
| 2N3R | | | | 0.7753 | 0.4341 | 0.0328 |
| 0N4R | | | | | 0.9982 | 0.5884 |
| 1N4R | | | | | | 0.8890 |

**Supplementary table 8.** Tukey-corrected post-hoc multiple comparisons for presynaptic area of 6A vulnerable neurons

| | 0N3R | 1N3R | 2N3R | 0N4R | 1N4R | 2N4R |
|---|---|---|---|---|---|---|
| Control | <.0001 | 0.0001 | 0.0019 | <.0001 | <.0001 | <.0001 |
| 0N3R | | 0.6000 | 0.1306 | 0.9930 | 0.9714 | 0.1323 |
| 1N3R | | | 0.9728 | 0.2259 | 0.1473 | 0.0010 |
| 2N3R | | | | 0.0268 | 0.0146 | <.0001 |
| 0N4R | | | | | 1.0000 | 0.4894 |
| 1N4R | | | | | | 0.6290 |

### Supplementary figures

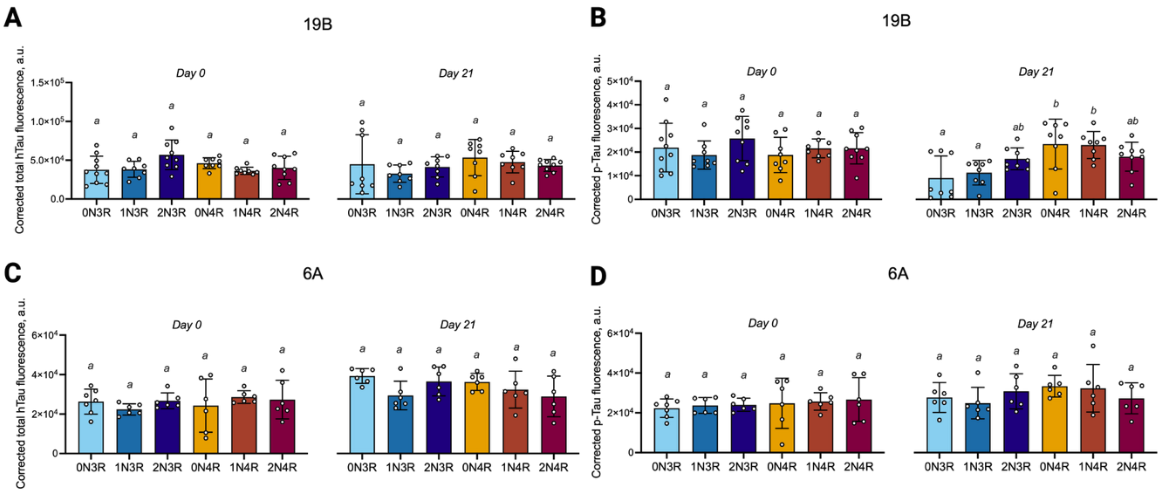


**Figure S1.** Quantification of total hTau and AT8 phosphorylation (p-Tau) levels in resilient (19B) and vulnerable (6A) neurons. The levels were quantified as corrected total cell fluorescence at day 0 and day 21 post-eclosion. Statistical significance was assessed using two-way ANOVA with post-hoc Tukey-corrected multiple comparisons; groups sharing a letter are not significantly different ($p < 0.05$). **A** Total hTau levels in resilient 19B neurons were broadly comparable across isoforms and did not increase with age. **B** p-Tau levels in the same neurons were similar across isoforms at day 0, but by day 21 isoform-specific differences emerged, with 0N4R and 1N4R showing higher p-Tau levels than 0N3R and 1N3R. **C, D** In vulnerable 6A neurons, total hTau and p-Tau levels both increased over time but remained broadly comparable across isoforms.
